# A spatial transcriptomics atlas of live donors reveals unique zonation patterns in the healthy human liver

**DOI:** 10.1101/2025.02.22.639181

**Authors:** Oran Yakubovsky, Amichay Afriat, Adi Egozi, Keren Bahar Halpern, Tal Barkai, Yotam Harnik, Yael Korem Kohanim, Roy Novoselsky, Ofra Golani, Inna Goliand, Yoseph Addadi, Merav Kedmi, Hadas Keren-Shaul, Liat Fellus-Alyagor, Dana Hirsch, Chen Mayer, Ron Pery, Niv Pencovich, Timucin Taner, Ido Nachmany, Shalev Itzkovitz

**Affiliations:** Department of Molecular Cell Biology, Weizmann Institute of Science, Rehovot, Israel; Department of General Surgery and Transplantation, Sheba Medical Center, Ramat Gan, Israel; Faculty of Medicine and Health Sciences, Tel Aviv University, Tel Aviv, Israel; Sheba Medical Center, Ramat Gan, Israel; Department of Surgery, Mayo Clinic, Rochester, MN, United States; Department of Immunology, Mayo Clinic, Rochester, MN, United States; Department of Immunobiology, Yale University School of Medicine, New Haven, CT, USA; Department of Life Sciences Core Facilities, Weizmann Institute of Science, Rehovot, Israel; Institute of Pathology, Sheba Medical Center, Ramat Gan, Israel; Lewis-Sigler Institute for Integrative Genomics, Princeton University, Princeton, NJ, USA; Department of Veterinary Resources, The Weizmann Institute of Science, Rehovot, Israel

## Abstract

Reconstructing gene expression atlases for human tissues is challenging due to limited access to healthy samples from live donors. Neurologically deceased donors often show ischemic changes, while tissues near diseased regions may have altered gene expression. The liver, with its unique regenerative capacity, allows analysis from live healthy donors (LHDs). Using spatial transcriptomics (Visum, Visium HD and MERFISH), we analyzed 16 liver samples: eight from young LHDs and eight from patients with liver pathology, sampling ‘adjacent normal’ tissue. LHD livers displayed significant gene expression differences from ‘adjacent normal’ tissues. Hepatocytes exhibited marked zonation along the porto-central axis of liver lobules, with key functions pericentrally shifted compared to other mammals. Our atlas identified dynamic programs in early steatotic hepatocytes, showing transitions from lipid uptake in low-lipid regions to insulin hypersensitivity in high-lipid regions. This study presents a spatial gene expression reference for the healthy human liver and insights into hepatocyte adaptations in steatosis.

## Main

The liver is a multitasking organ, implementing diverse functions ranging from maintenance of nutrient homeostasis, circulating protein production, bile acid secretion and foreign substance detoxification^1^. Hepatocytes, the parenchymal liver cells that perform these tasks, operate in structured anatomical units called liver lobules. These hexagonal units, measuring 0.5-1mm in diameter in mammals, are polarized by blood that flows from portal veins and drains into central veins. This polarized blood flow generates gradients of oxygen, nutrients and morphogens. As a potential adaptation to this graded microenvironment, key hepatic functions are expressed in distinct lobule layers, a phenomenon termed ‘Liver zonation’ ^1,2^. Many liver diseases, including metabolic-associated steatotic liver disease^3^(MASLD), autoimmune hepatitis^4^, cholangitis^5^ and parasite infections^6,7^ show zonated pathological patterns. Detailed spatial expression maps of hepatocytes in humans are therefore critical as a reference to understand human liver pathologies. In recent years, several cell atlases of the human liver have been reconstructed, yielding important insights into its cellular states^8–17^. Notably, challenges with tissue acquisition impeded our ability to analyze human liver samples from completely healthy individuals.

The selection of the most appropriate healthy reference tissue for human cell atlas studies is a critical consideration^18^. Human liver cell atlases have thus far been reconstructed from either neurologically deceased liver donors (NDD)^9,19^ or from histologically-healthy appearing tissues adjacent to a liver pathology that warranted surgical intervention^11,15^. Although these samples are considered healthy liver tissues, several factors challenge their validity as a reference point to the healthy liver^20,21^. Livers from neurologically deceased donors undergo metabolic disturbances resulting from long-term positive mechanical ventilation, non-enteral nutrition, and medication influences, which can all alter hepatic gene expression in the pre-liver-extraction period. Moreover, post-mortem tissues are subject to variable ischemia times, leading to non-random and transcript-dependent RNA degradation; indeed, ischemia-dependent RNA degradation can account for up to 40% of the variability in RNA-seq tissue studies^21^. Liver samples that are obtained from regions adjacent to liver lesions may themselves be impacted by the disease. Indeed, liver gene expression can change even in patients with extra-hepatic tumors^22^. Here, to establish a ‘true-healthy’ human liver spatial transcriptomics atlas devoid of any perturbations or pathological conditions, we analyzed samples from LHDs. These individuals undergo comprehensive medical screening to ensure a healthy liver physiology baseline, enabling them to donate a significant portion of their liver to a designated recipient.

We employed 10x Visium spatial transcriptomics and MERFISH^23^ to reconstruct liver zonation in LHDs and compared the results to the ‘adjacent-normal’ samples, identifying profound expression differences. We also reconstructed spatially resolved atlases of other mammalian species to identify unique and shared zonation features with humans. Given that LHD liver transplants are deemed suitable for transplantation with up to 20% steatosis, we were able to analyze a spectrum of steatosis levels on the background of otherwise completely healthy livers. We integrated spatial transcriptomics with machine learning-based pixel classification of the underlying H&E images to stratify zonal hepatocytes by lipid content and reconstruct their dynamic changes in gene expression along the course of lipid accumulation. Our study provides a resource for understanding the human liver and highlights spatial-dependent hepatocyte adaptations in the early stages of steatosis.

## Results

### LHD liver tissues are transcriptomically unique

To assemble a detailed gene expression atlas of the human liver, we collected liver specimens from two procedure types: the first is live organ donor hepatectomy, in which a portion of the liver is surgically removed from an LHD before it is transplanted in the recipient. The second is liver resection, in which a portion of the liver is resected due to a liver pathology (Fig. 1a, Methods). We also performed spatial transcriptomics of livers from other mammals to identify shared and unique zonated expression patterns (Fig. 1a). Adjacent normal samples exhibited considerable heterogeneity in age, BMI, smoking status and prior medications (Fig. 1b). In contrast, the LHD samples demonstrated remarkable homogeneity in these features. We categorized the samples according to their histological diagnosis. Our cohort consisted of 8 LHDs, 3 of which exhibited varying degrees of mild steatosis and 8 adjacent normal sample donors, 5 of which exhibited varying degrees of steatosis, 2 with portal fibrosis, and 1 patient who underwent Associating Liver Partition and Portal vein Ligation for Staged hepatectomy^13^ (ALPPS) (Fig. 1a-b). We performed 10X Visium spatial transcriptomics on fresh frozen samples, obtaining the transcriptomic signatures of 49,092 spots, with a median of 3,008 spots per patient and 3,393 Unique Molecular Identifiers (UMIs) per spot (Extended Data Fig. 1). We filtered outlier spots and retained 44,243 high-quality spots (Methods).

**Figure 1.**
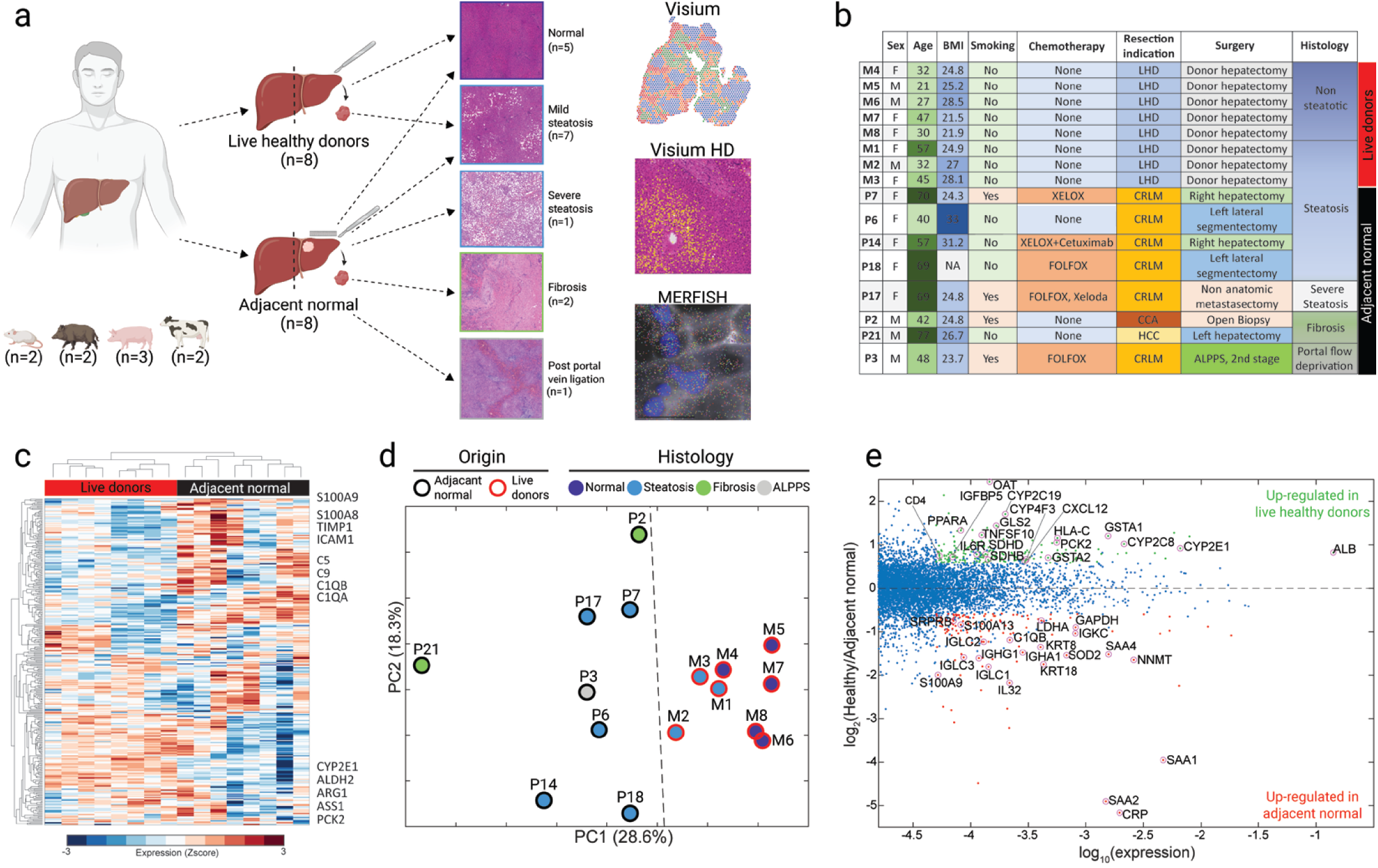
A spatial expression atlas of human and mammalian livers. **a.** Study design. Liver tissue gene expression was measured using 10X Visium spatial transcriptomics (human n = 16, mouse n=2, boar n=2, domesticated pig n=3, cow n=2), Visium HD (n=2) and MERFISH (human n=2). Diagram created using BioRender. **b.** Clinical characteristics of patients from whom liver samples were collected (Methods). CRLM, colorectal liver metastases; CCA, cholangiocarcinoma; HCC, hepatocellular carcinoma. **c.** Heatmap of hierarchical clustering of pseudo-bulk transcriptomic profiles. Analysis done on highly expressed genes. Representative genes are shown on the right. Expression units are Z-scores of log10 UMI-sum normalized expression. **d.** Principal component analysis of pseudo-bulk transcriptomic profiles. LHD is in red outline, and adjacent normal livers are in black outline. Dot colors denote pathological classification - no pathology (dark blue), steatosis (light blue), fibrosis (green), and portal vein ligation (grey). Panels **c,d** based on 323 genes with a maximal expression across patients above 5e-4 (Methods). **e.** MA plot showing the gene expression ratios between the LHD samples and the adjacent normal samples. Genes with maximal expression above 5e-5, q-value below 0.25, and fold change ratio above 1.5 are highlighted in color. Genes upregulated in the LHD are in green, genes upregulated in the adjacent normal are in red. Q-values calculated using the Benjamini-Hochberg procedure. Rank-sum p-values were calculated for all genes that had maximal expression above 5e-5 in all the 16 patients. Selected markers for distinct immune functions and classic hepatocyte functional genes are highlighted with a circle.

Pseudo-bulk and differential gene expression revealed distinct sample clustering based on clinical context, particularly distinguishing LHDs from adjacent normal livers (Fig. 1c-e, Supplementary Table 1). Adjacent normal livers exhibited higher expression of immune-related genes. These included the calprotectin genes *S100A8* and *S100A9*, the Kupffer cell markers *C1QA* and *C1QB*, and the plasma cells genes *IGHA1* and *IGHG3*. Adjacent normal livers further showed elevated levels of the serum amyloid genes *SAA1* and *SAA2*, the complement system activator *CRP* and the matrix metalloproteinase *TIMP1.* In contrast, LHD livers exhibited elevated expression levels of classic hepatocyte functional genes, such as the urea cycle enzymes *ASS1*, cytochrome P450 enzymes *CYP2E1* and *CYP2C8*, and the gluconeogenic gene *PCK2* (Fig. 1c, e). We further analyzed the liver expression programs of an NDD^9^ (Extended Data Fig. 1e, Methods). NDD liver also exhibited elevated expression of stress-related programs (IFITM2,3, SAA1,2), as well as elevation of the hypoxia induced genes LDHA and GAPDH. NDDs showed lower expression levels of some hepatocyte genes (ALB, CYP2E1), yet comparable levels of others (ASS1, Extended Data Fig. 1e). Our analysis, therefore, underscores the expression differences between LHD livers and adjacent normal and NDD livers. We next pursued the analyses of LHD livers to establish the zonation patterns of the ‘true healthy’ human liver.

### Zonation patterns in the healthy human liver

To assemble a spatial reference frame, we determined the location of each spot along the porto-central axis using the gene expression levels of zonated landmark genes. To this end, we validated the consistent pericentral expression of *CYP2E1*^24,25^ in hepatocytes close/far from portal triads, as labeled by a certified pathologist (Extended Data Fig. 2a). Next, we extracted the 20 genes that were most correlated or anticorrelated with *CYP2E1* among spots and defined them as our pericentral and periportal landmark gene sets. Those landmark gene sets’ normalized summed expression levels enabled assigning a percentile-categorized zone index to each spot (Fig. 2a, Methods). We grouped spots by their lobule zones and averaged the expression of all spots from the same zone. Out of 1,724 genes that were identified as hepatocyte-specific genes (Methods), 1,141 exhibited expression levels that were significantly different across zones (Fig. 2b-c, q-value<0.25), demonstrating that most human hepatocyte genes are zonated (Supplementary Table 2). To validate our reconstructed zonation profiles, we applied multiplexed single molecule transcript imaging of 500 genes using the MERSCOPE platform (Extended Data Fig. 2b,c, Methods). Zonation profiles strongly overlapped between the MERFISH and Visium measurements (n=2, spearman correlation: patient M5: R=0.49 p=7.4e-7, patient M8: R=0.57 p<5e-324, Extended Data Fig. 2d-e, Methods). To achieve higher spatial resolution, we also applied high-definition Visium HD measurements of two patient sections (Extended Data Fig. 3). One of the samples (M6) yielded high RNA capture rates, enabling robust zonation reconstruction with a strong overlap to the zonation profiles obtained with the lower resolution Visium slides (Extended Data Fig. 3). The zonation profiles in the healthy human liver can be explored using our online web app (<u>link</u>).

**Figure 2.**
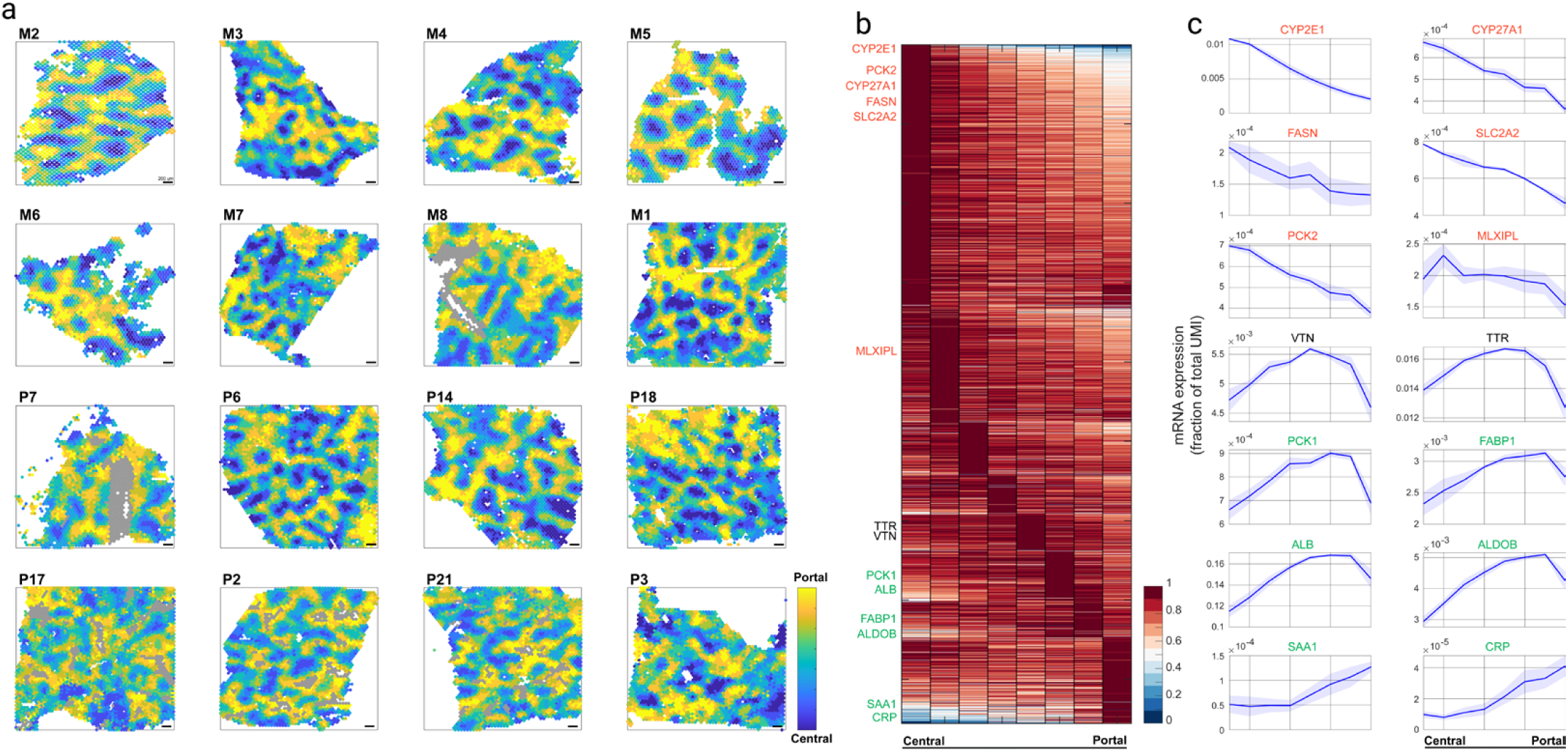
Zonation of hepatocyte gene expression. **a.** Spatial transcriptomics Visium 10x liver slides, colored by spot-inferred zone index (Methods) in a range from Yellow (portal) to blue (central). Spots predominantly composed of fibrotic tissue are represented in gray (Methods). **b.** Heat map of the mRNA zonation profiles based on the Visium 10X data. In total, 1,711 highly expressed hepatocyte-specific genes with a gene zonation score (Methods) are shown. Profiles are normalized to their maximal value across zones across all healthy, non-steatotic liver samples (M4-M8, fig 1b). Profiles are sorted by the zonation profiles centers of mass (Methods). **c.** mRNA zonation profiles of representative hepatocyte-specific genes in (b). Data are means (lines) and standard errors of the means (SEM, patches) across all healthy, non-steatotic liver samples’ zonation profiles.

We found that pericentral hepatocytes showed elevated levels of the xenobiotic metabolism genes such as *CYP2E1*, bile acid biosynthesis genes such as *CYP27A1*, and lipid biosynthesis genes such as *FASN* and *MLXIPL*. Periportal hepatocytes showed elevated expression of immune-related genes, such as *CRP* and *SAA1*, as well as the gluconeogenic genes *PCK1* and *ALDOB*. Notably, the mitochondrial gluconeogenesis gene *PCK2*, exclusively expressed in humans, showed strong pericentral zonation. This pattern contrasts with the view that gluconeogenesis is an exclusively periportal function in mammals^24,26^. The main hepatic glucose transporter GLUT2, encoded by the gene *SLC2A2* was also pericentrally zonated (Fig. 2b,c).

We used gene set enrichment analysis^27^ (GSEA) to identify extensive pericentral enrichment of multiple hepatic functions, including xenobiotic metabolism, fatty acid metabolism, and peroxisome genes (Extended Data Fig. 4a). We also identified zonation of ligands^28^ (Extended Data Fig. 4b) and receptors^28^ (Extended Data Fig. 4c). Zonated examples included the pericentral Wnt ligands (*WNT2*, *RSPO3*) and the *RSPO3* receptor *LGR5*^29–31^, as well as the periportal Notch ligands (*JAG1,2*) and receptor (*NOTCH3*). Zonated transcription factors^32^ (TFs, Extended Data Fig. 4d) included the pericentral aryl hydrocarbon receptor *AHR*, an activator of P450 xenobiotic metabolism genes^33^ and the periportal Notch activated TFs *HES1*, *HEY1*, *HEYL*. Notably, *HNF4A*, a key hepatic transcription factor that is more highly active in periportal hepatocytes in mice^26^ showed strong pericentral zonation in humans (Extended Data Fig. 4d, q-value=8.5e-14).

### Pericentral shifts in human hepatocytes compared to other mammals

Our zonation analysis indicated that key hepatocyte functions and TFs are pericentrally zonated. To examine whether this phenomenon is distinct to humans, we compared zonation profiles to those in other mammals. To this end, we analyzed a published dataset of mice spatial transcriptomics^34^ (n=2, Fig. 3a), and assembled a new dataset of spatial transcriptomics of 3 additional mammalian species with body sizes and metabolic rates more comparable to human: wild boar (n=2, Fig. 3b), cow (n=2, Fig. 3c), and domestic pig (n=3, Fig. 3d). We performed 10X Visium spatial transcriptomics on fresh frozen samples (Methods, Fig. 3b-d, Extended Data Fig. 5a-d). We determined each spot’s zone based on landmark gene expression in cow and mouse and based on distances from the portal fibrotic septa in pigs and boars (Methods). For hepatocyte genes with 1:1 orthologs across all species (n=1,344 genes, Methods), we computed the log2 ratio between the periportal and pericentral expression levels, normalized to hepatocyte genes. We found that hepatic genes were significantly biased toward the pericentral zone in humans (Fig. 3e, one-tailed rank-sum p<3.4e-15 for all species). Portal hepatocytes have been shown to specialize in the translation of hepatic-secreted proteins in mice^24,26^. This zonal trend was suggested to be optimal as it co-localizes one of the most energetically demanding cellular tasks of protein translation to the highly oxygenated zone in the tissue, where ATP could be more readily produced by respiration. Notably, we found that in humans, secreted proteins were also significantly shifted pericentrally (Extended Data Fig. 5e, one-tailed rank-sum p<5.22e-3), except for complement system proteins (Extended Data Fig. 4f, one-tailed rank-sum, p=0.06). Indeed, we found that respiration genes in humans also showed a significant pericentral shift compared to mice and other species (Fig. 3f). The higher expression of respiration genes in pericentral human hepatocytes could potentially supply the energy needed for protein secretion in this zone.

**Figure 3.**
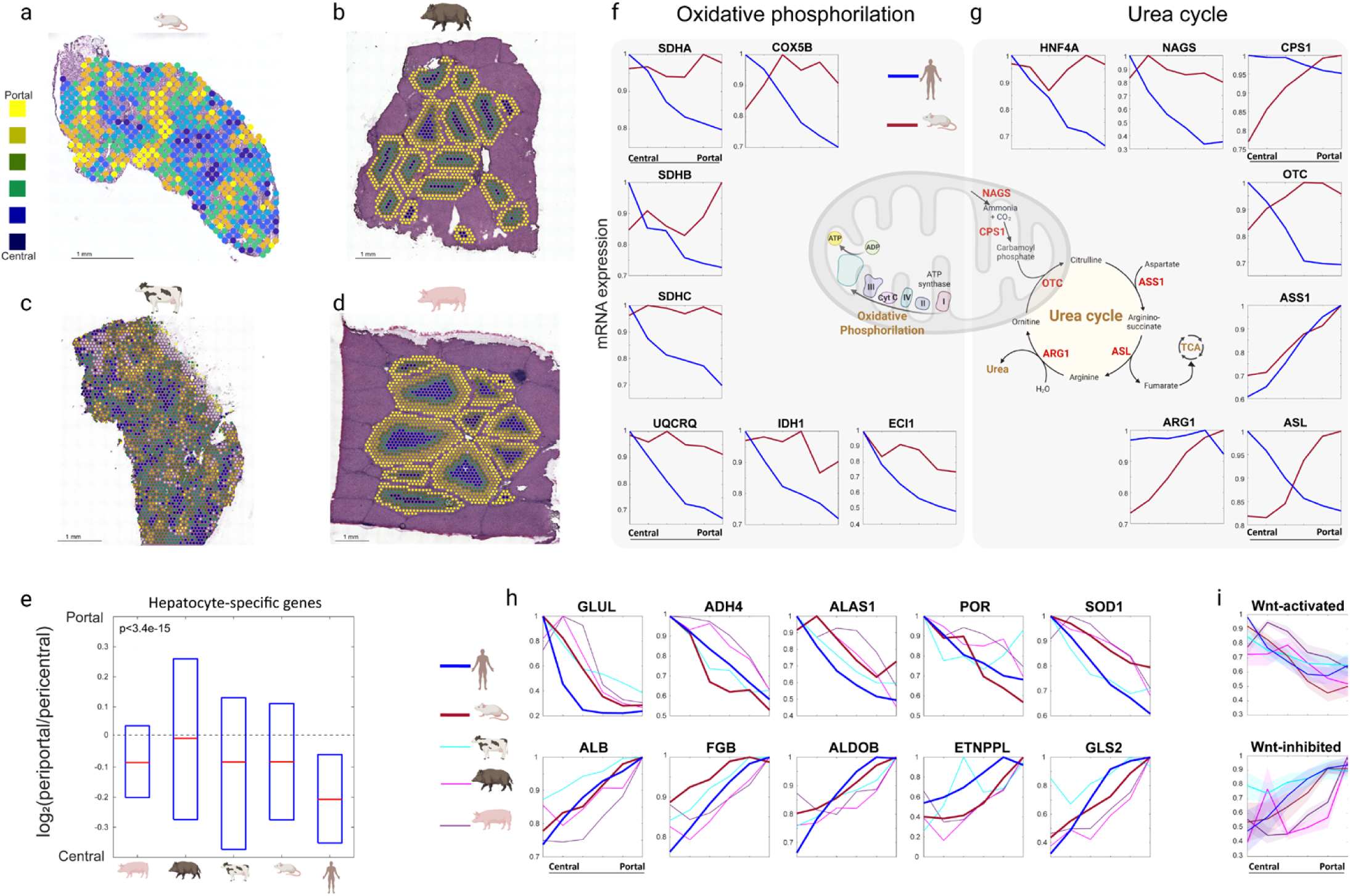
Distinct and shared zonation programs across mammals. a-d. Representative spatial transcriptomics Visium 10x liver slides, colored by spot-inferred zone index (Methods) in a range from Yellow (portal) to blue (central). **e.** Boxplot showing the portal bias 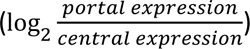 of highly expressed hepatocyte-specific genes (Methods). P-value is the minimal among the ranksum test p-values between human and each of the other four species individually. Red lines are medians, boxes span the interquartile range. **f-g.** mRNA zonation profiles of highly expressed key genes in oxidative phosphorylation pathway in (f), and urea cycle in (g). **h.** Zonation profiles of representative genes with consistent zonation patterns across the mammals studied. Lines in **f-h** are means over the individual repeats within each species, normalized to the maximum across zones. **i**. Cross-species zonation profiles of Wnt-activated (upper panel) and Wnt-inhibited (lower panel) genes. Lines/patches are relative means/SEMs over the max-normalized zonation profiles of the genes within each set. Gene sets are the most Wnt-up-regulated and Wnt-down-regulated genes analyzed in Gougelet et al (Methods)^39^. Profiles are normalized to their maximal expression across zones. blue-human, red-murine, cyan-cow, pink-boar, purple-domesticated pig. Human n = 7, mouse n=2, cow n=2, boar n=2, domesticated pig n=3.

The urea cycle is an important hepatic process by which toxic ammonia is converted into urea that is excreted in the urine. The cycle contains multiple mitochondrial and cytosolic enzymes (Fig. 3g). We found that, except for the periportal enzyme *ASS1*, the expression of all the urea cycle enzymes was shifted pericentrally in humans compared to mice (Fig. 3g). The pericentral expression of *OTC* could be related to the pericentral zonation of *HNF4a*, an up-stream regulator of this gene^35^ (Fig. 3g). *ASL*, one of the key enzymes in the urea cycle, produces both arginine, which feeds the urea cycle, as well as fumarate, which can feed the TCA cycle. The pericentral zonation of *ASL* in humans, distinct from that in mice, could also serve to increase the respiratory activity of pericentral hepatocytes.

In contrast to the pronounced distinct pericentral zonation of key hepatocyte genes in humans, we also found universally-zonated genes that showed similar trends across organisms. These included the pericentral *GLUL* and *ADH4*, and the periportal *ALB* and *GLS2* (Fig. 3h). *GLS2* encodes glutaminase, which breaks down glutamine into glutamate, whereas *GLUL* encodes glutamine synthetase, which re-synthesizes glutamine from glutamate. The periportal expression of *GLS2* and pericentral expression of *GLUL* matches the need to recycle this amino acid to ensure balanced representation in the liver entry and exit^26^.

The Wnt pathway constitutes a major regulator of liver zonation in mice^36,37,37–41^. Our ligand analysis indicated that the key Wnt ligands *WNT2* and *RSPO3* are pericentrally zonated in humans (Extended Data Fig. 4b-d). To examine the overall zonation of Wnt pathway genes we reconstructed the mean zonation profiles of Wnt-activated and inhibited genes^24^. We found that Wnt-activated/inhibited genes were pericentrally/periportally zonated respectively in all analyzed species (Fig. 3i, Extended Data Fig. 6). Wnt pathway therefore seems to be an evolutionary conserved regulator of liver zonation.

### Functional changes along the course of steatosis

Metabolic associated steatotic liver disease consists of abnormal fat accumulation in hepatocytes. The disease impacts 25% of the population^42,43^ and consists of stages defined by increasing hepatic lipid content and eventual inflammation and fibrosis^44,45^. Hepatic lipid accumulation can be driven by multiple processes, ranging from elevated plasma levels of fatty acids, increased fatty acid import or synthesis, or decreased lipid export and breakdown. MASLD has a strong zonal feature, and lipid accumulation in adults largely commences in the pericentral zone^46,47^. Our cohort included 4 patients with different levels of lipid accumulation (M1, M2, M3, P6, Fig. 4a-d). We applied machine learning algorithms to quantify the amounts of lipids per spot (Methods, Fig. 4a-c, Supplementary Table 6). Our patients exhibited pronounced pericentral bias in lipid content (Extended Data Fig. 7). Our spatially resolved atlas enabled exploring functional changes in hepatocytes at increasing lipid levels. To this end, we ranked pericentral spots by their binned lipid contents and averaged gene expression across ranges of hepatic lipid accumulation (Fig. 4e-k, Supplementary Table 3).

**Figure 4.**
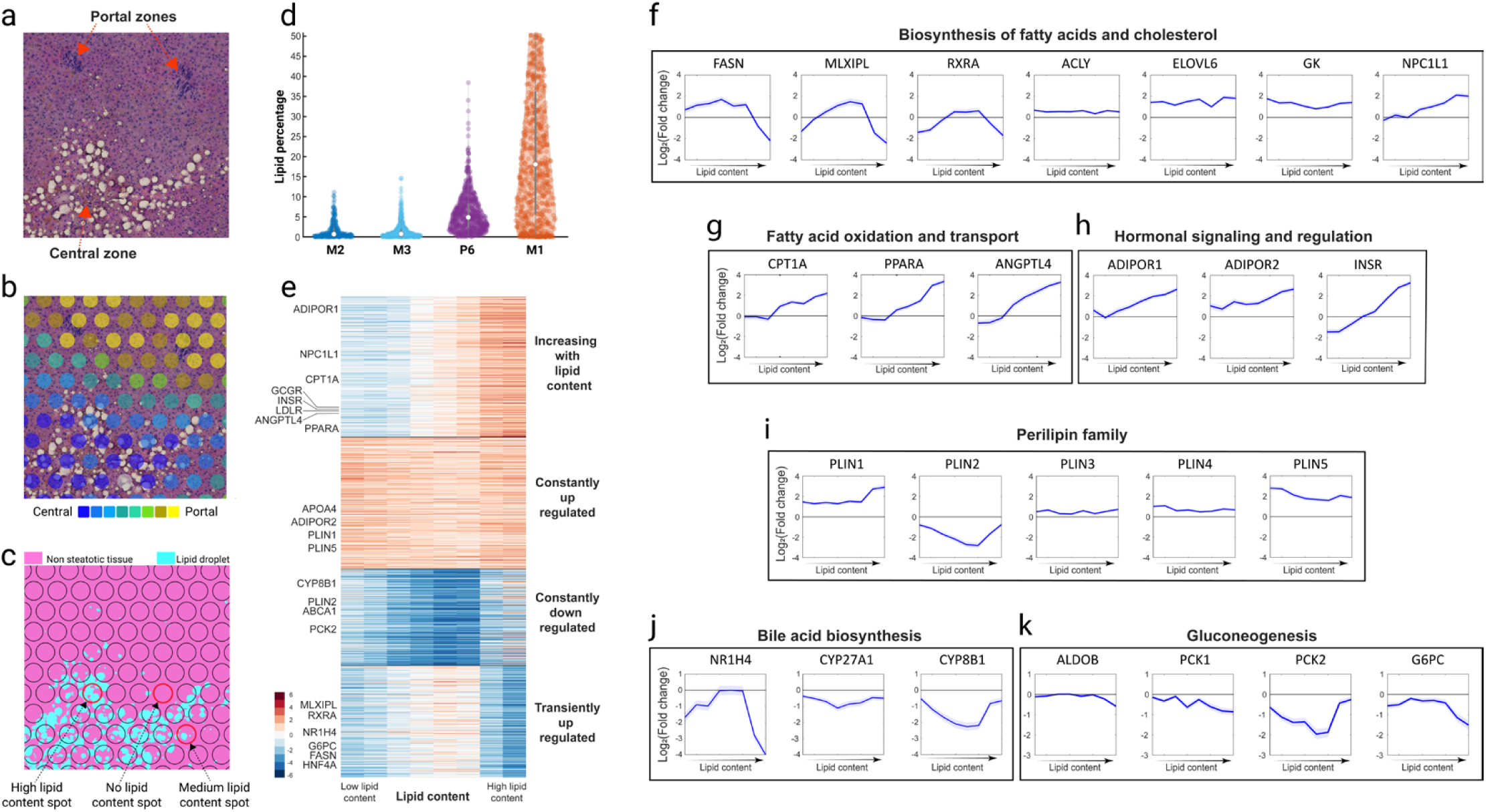
dynamic gene expression changes in steatotic hepatocytes. **a.** H&E-stained Visium slide demonstrating the centrally biased zonation of lipid droplets (sample M1). **b.** Overlayed spots colored by their inferred zone index for the same image in (a). **c.** Overlayed color-coded pixel classification results for the same image as in (a) and (b). color codes: Non-steatotic tissue-pink, lipid droplets-cyan. **d.** Distributions of lipid areas in pericentral spots (n=4) according to the pixel-classification of (c). **e.** K-means clustering (K=4) of the highly expressed significant genes based on Log2 of fold change of each gene in the steatotic spots over the non-steatotic median expression. Included are genes with maximal expression above 5e-6 and q-value<0.25. **f-k.** Examples of expression changes across lipid content bins for representative genes grouped by functional classes. Lines are means over the spots in each lipid content bin, patches are SEMs.

Our analysis revealed a bi-phasic behavior of hepatocytes, which shifted from preferential lipid accumulation to adaptations that avoid hepatic steatosis. *FASN*, encoding fatty acid synthase^48^, its activator *RXRA*^49^, and the transcription factor CHREBP (encoded by *MLXIPL*), which controls glucose conversion to acetyl-coA for fatty acid biosynthesis^48^, exhibited an initial increase with hepatic lipid content followed by a decline at higher lipid content (Fig. 4f). Conversely, *CPT1A* and *PPARA*, central genes in the machinery for breakdown of intra-hepatic lipids through fatty acid oxidation, were elevated as lipid content increased. Hepatocytes with elevated lipid content also up-regulated *ANGPTL4*, a liver-secreted hormone that inhibits the release of circulating fatty acids from adipocytes^50^(Fig. 4g). This could potentially be a systemic cue for the need to lower circulating fatty acids upon physiological saturation of the internal lipid accumulation capacity of hepatocytes. High lipid content was also associated with an increase in the insulin receptor *INSR* and in the receptors for adiponectin (*ADIPOR1*, *ADIPOR2*, Fig. 4h), a hormone that increases insulin sensitivity. These trends could potentially be an adaptation for hepatocyte insulin resistance at high lipid content^51^. Additional genes involved in lipid biosynthesis such as ATP citrate lyase (*ACLY*), which converts citrate to Acetyl-CoA and the fatty-acid elongase *ELOVL6* showed elevation compared to non-steatotic hepatocytes regardless of lipid content (Fig. 4f). Perilipin genes, which generally promote hepatic lipid accumulation, showed elevated levels, except for *PLIN2*, which showed reduced levels in steatotic hepatocytes (Fig. 4i). Hepatocyte genes encoding other homeostatic processes, such as bile acid biosynthesis and gluconeogenesis were down-regulated throughout the lipid-accumulation process (Fig. 4j,k). Our analysis of hepatic gene expression changes along the course of steatosis can be explored using our online web app (<u>link</u>).

## Discussion

Our study presents the spatial expression blueprint of the healthy human liver and reveals unique hepatocyte zonation features that are distinct from other mammals. Specifically, we found that pericentral hepatocytes in humans engage in diverse hepatic functions, including ones that are preferentially performed by portal hepatocytes in other mammals. These include energetically demanding tasks such as protein secretion, some gluconeogenic components (*PCK2*) and key enzymes in the urea cycle (*ASL*, *NAGS*, *OTC*). Moreover, we identified that *HNF4*, a key hepatic transcription factor which was shown to be periportally-zonated in mice, is significantly pericentrally-zonated in humans (Fig. 3g). The distinctly pericentral functions in humans compared to other mammals included cellular respiration. This elevation might facilitate the needed energy for the diverse functions employed in the pericentral zone. An additional gene that was pericentrally shifted in humans compared to mice was *SLC2A2*, encoding the glucose transporter GLUT2, (Fig. 2c, Supplementary Table 2). The higher expression of GLUT2 in pericentral hepatocytes may be in line with the ‘glucose paradox’ observation that hepatic glycogen stores are replenished directly from glucose in pericentral hepatocytes, yet from lactate trough gluconeogenesis in periportal hepatocytes^52^. Indeed, our analysis also revealed a higher periportal expression of lactate dehydrogenase b (*LDHB*), the enzyme that converts lactate into pyruvate, yet a non-zonated expression of lactate dehydrogenase a (*LDHA*) that converts pyruvate to lactate (Supplementary Table 2).

Our spatially resolved expression atlas enabled analysis of hepatic gene expression programs along the course of early steatosis, in the background of healthy individuals. At these stages, before the pathologic commencement of inflammation or fibrosis, we found that hepatocytes seem to be initially tuned towards lipid accumulation. This could be a physiological adaptation to reduce the systemic impact of circulating free fatty acids. Adaptations to lipid accumulation involve multiple organs and require systemic hormonal changes, partially driven by insulin resistance^51^. Notably, we found that pericentral steatotic hepatocytes elevate the sensing machinery for insulin, including *INSR*, *ADIPOR1* and *ADIPOR2*, potentially adapting to developing cell-intrinsic insulin resistance. Notably, our analysis of steatosis focused on relatively early stages, before the commencement of hepatic inflammation (steato-hepatitis), a condition that has been explored in depth in several recent publications^11,53^.

Our combined analysis of spatial transcriptomics and lipid content, quantified based on machine learning of H&E images, could be applied to other organs. For example, it could be used to explore expression differences in adipocytes depending on their lipid content. Our spatial expression atlas forms a blueprint of the healthy liver state, which will form a reference for detailed analysis of zonal changes in diverse liver pathologies such as viral hepatitis, fibrosis, drug-induced liver injuries and auto-immune hepatitis.

## Supplementary information

**Extended Data Figure 1.**
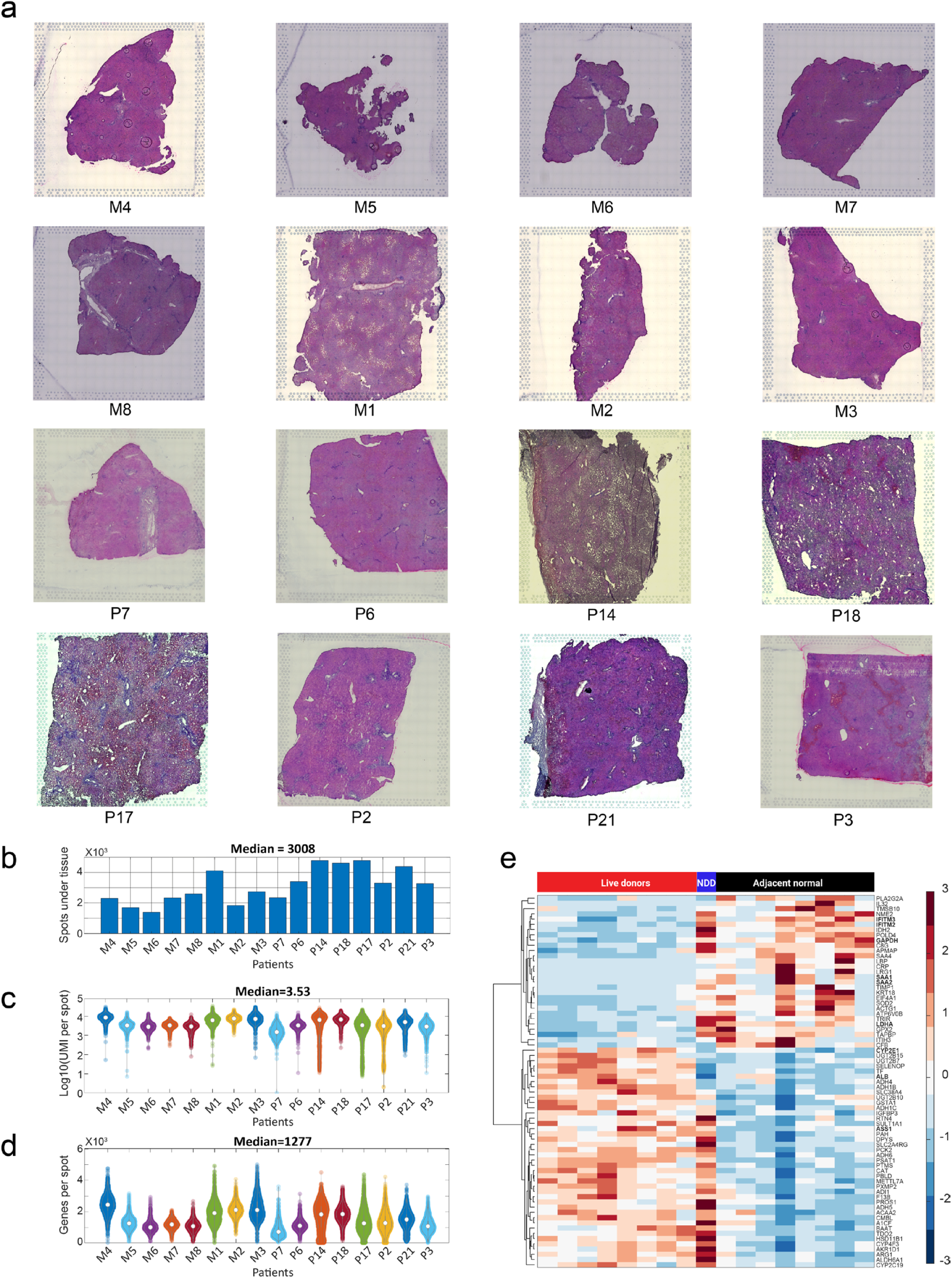
Images and quality control metrics for the sequenced human Visium samples. **a.** H&E images of tissues from Visium experiments. The capture area within the fiducial frame is 6.5×6.5 mm. Samples labeled with ‘M’ correspond to liver tissues obtained from LHDs. Samples labeled with ‘P’ correspond to adjacent normal liver tissues resected due to underlying pathological conditions. **b-d**. Quality metrics of Visium data, n = 16 patients examined in 4 independent experiments. Patient order matches that in Figure 1b. Circles in c-d are medians, gray box span interquartile ranges. **e.** Heatmap of hierarchical clustering of pseudo-bulk transcriptomic profiles of LHD, adjacent normal and NDD (Visium data from Andrews et al.^9^). Analysis done on genes showing differential expression between LHDs and adjacent normal livers (expression>2e-4, qval<0.05 and ratio above 1.5). Genes mentioned in the main text are in bold. Expression units are Z-scores of log10 UMI-sum normalized expression.

**Extended Data Figure 2.**
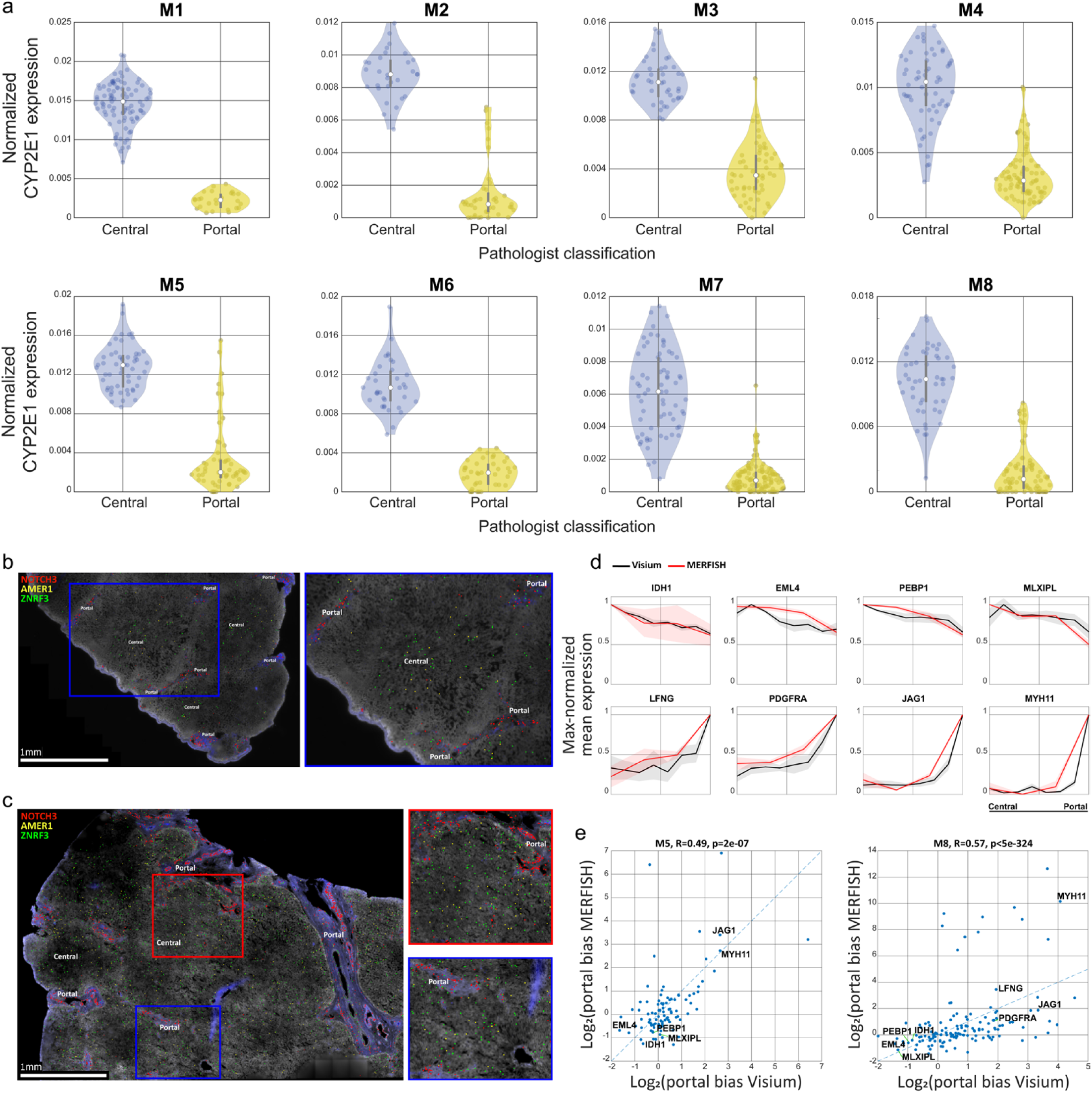
In-situ validation using MERFISH for zonation profiles computed from spatial transcriptomics. **a.** Violin plots showing the spatial distribution of *CYP2E1* across pericentral and periportal spots labeled by a certified pathologist. Kruskal-Wallis p<1.94e-12 for all samples. Circles are medians, gray boxes span interquartile ranges. **b-c.** MERFISH image of patient M5 (b) and M8 (c) showing selected zonated genes. Blowouts of selected regions are shown on the right. **d.** mRNA zonation profiles for representative genes. Profiles are means over the 2 patients in MERFISH (red, M5 and M8) and 5 patients in Visium (black, M4, M5, M6, M7, M8), normalized to their maximal expression level across zones, patches show the SEM. **e.** Spearman correlation between the portal bias 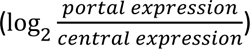 in the MERFISH and Visium methods. Shown are highly expressed genes (maximal expression above 5e-5). Selected genes’ names are shown.

**Extended Data Figure 3.**
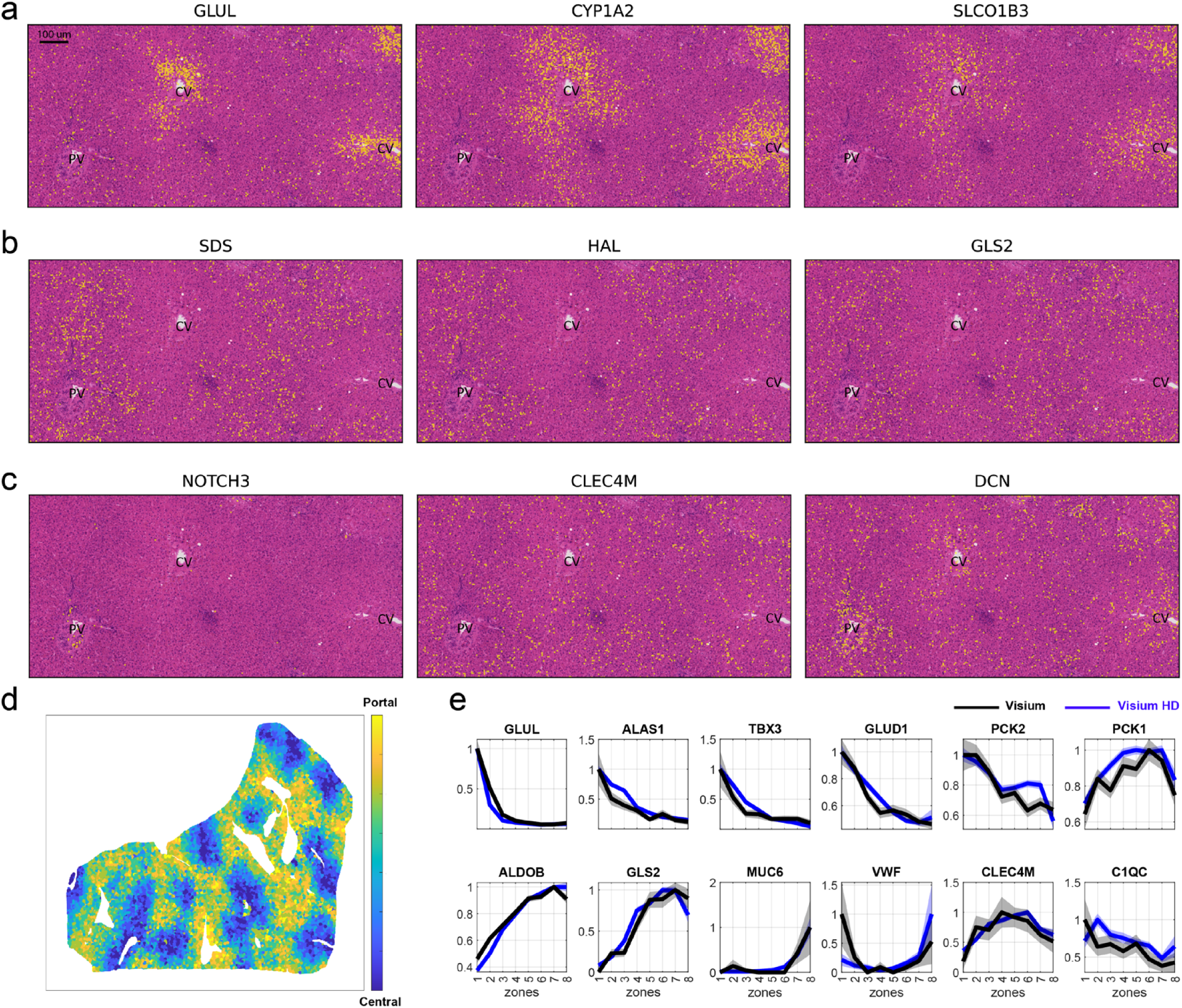
Visium HD provides zonation reconstruction with high spatial resolution. (patient M6)**. a-c.** Representative images of pericentral genes (a), periportal genes (b) and non-parenchymal genes (c) including the vascular smooth muscle cell marker NOTCH3, the sinusoidal endothelial cell marker CLEC4M and the fibroblast marker DCN. PV – portal vein, CV – central vein. **d.** Spatial map of 8*8 spots colored by inferred zone. **e.** Reconstructed zonation profiles for M6 based on classic Visium (black) and Visium HD (blue). Lines are means, patches are SEMs. Profiles normalized to their maximal expression across zones.

**Extended Data Figure 4.**
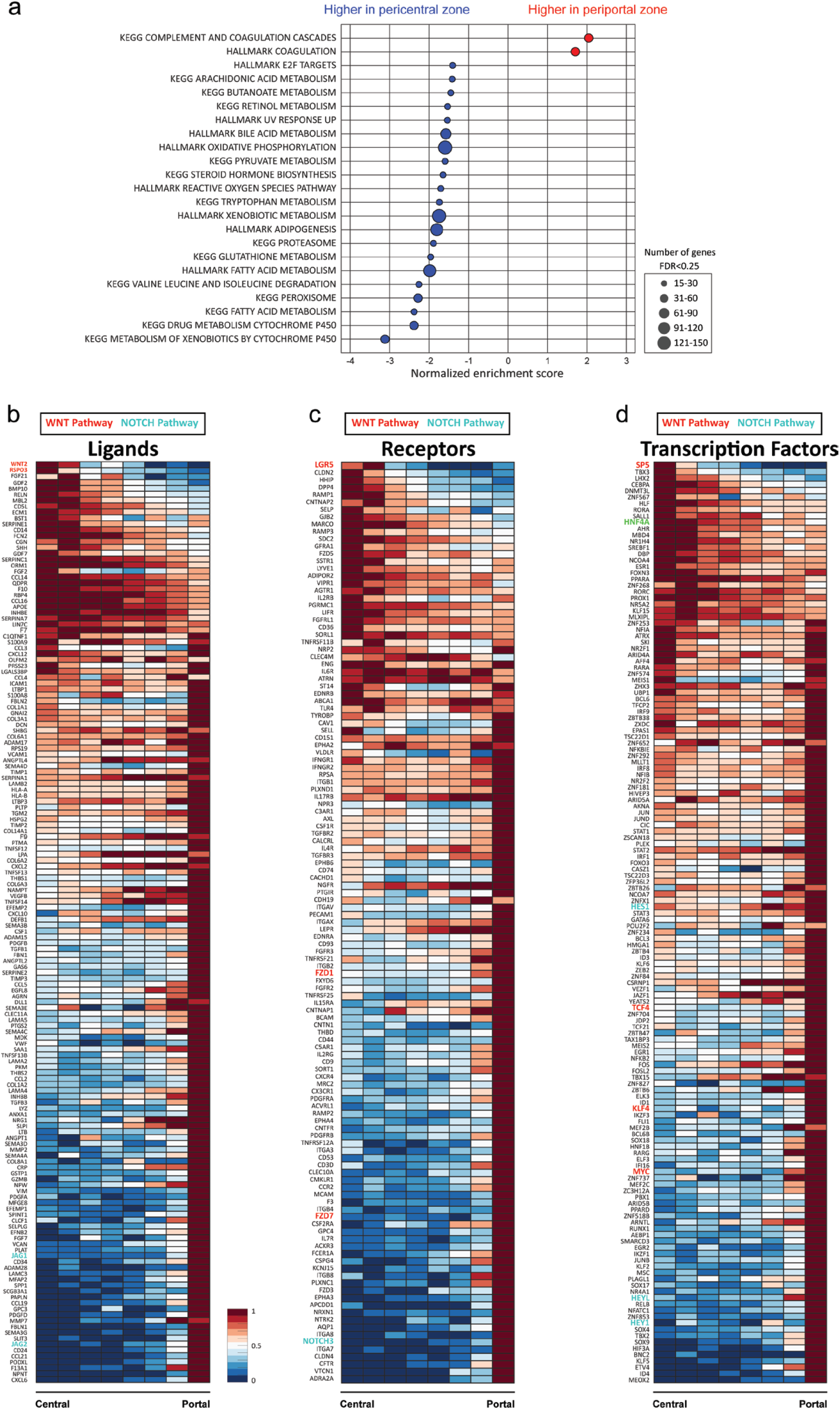
Pathway enrichment gene zonation analysis in healthy LHD liver samples. **a.** Gene Set Enrichment Analysis (GSEA)^27^ of the sorted gene zonation scores, calculated as the linear combination of the zone index with the highest gene expression and the gene’s center of mass. The analysis utilized the KEGG^54^ and Hallmark^55^ gene sets in healthy, non-steatotic LHD liver samples (n=5). **b-d.** Zonation patterns of ligands (b), receptors (c), and transcription factors (d) for highly expressed highly variable genes (q-value less than 0.05, expression more than 3e-6, dynamic range (ratio of maximal to minimal zonation value) more than 0.5) were analyzed in healthy, non-steatotic LHD liver samples. The cumulative significance of gene zonation across all patients (n = 5) was determined using Fisher’s method, combining the zonation bias p-values of each gene into a chi-squared distribution. Profiles are normalized by their maximum and sorted by centers of mass.

**Extended Data Figure 5.**
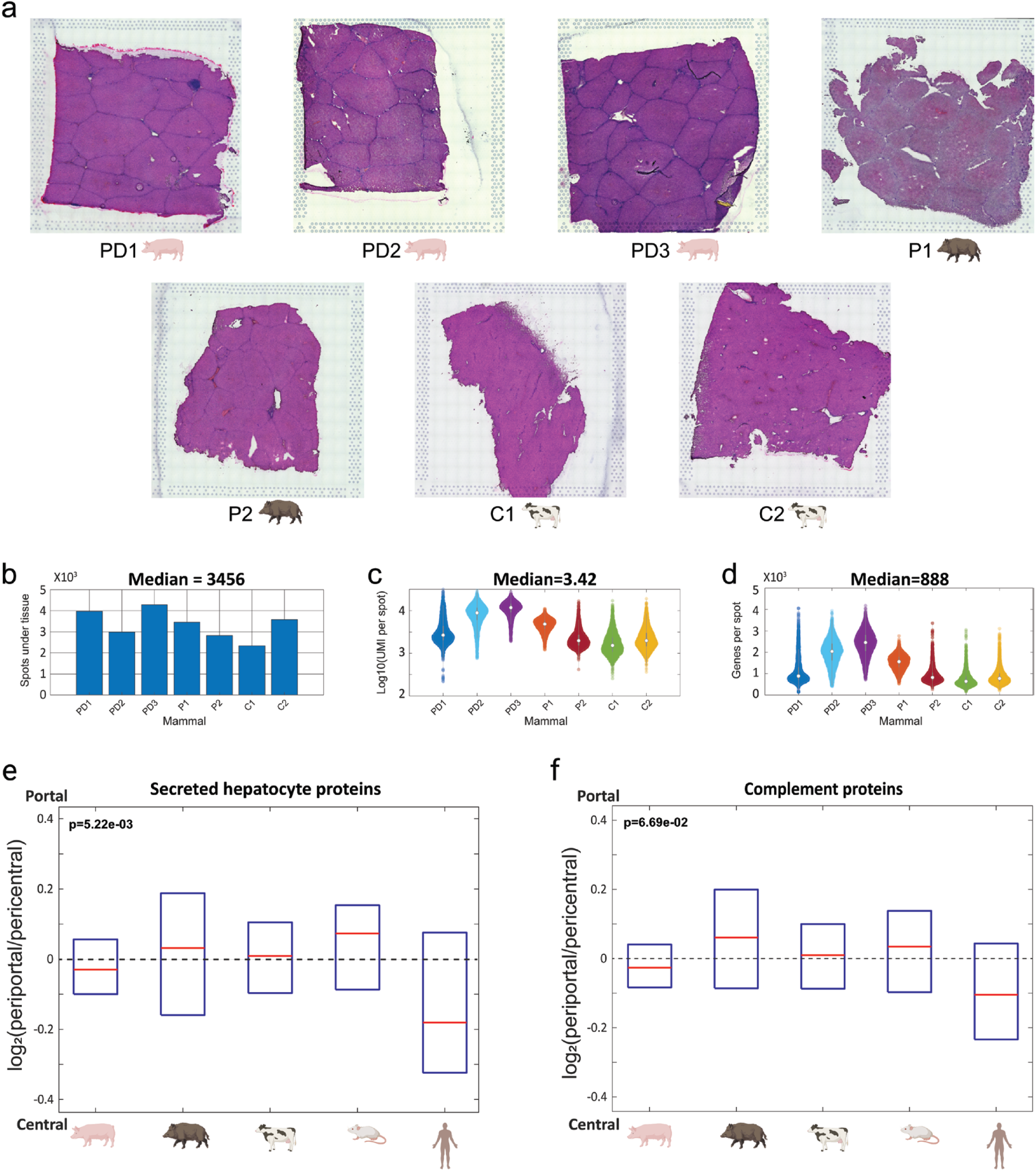
Non-human mammalian Visium samples. **a.** H&E images of tissues from spatial transcriptomics experiments. The capture area within the fiducial frame is 6.5 ×6.5 mm. ‘C’ - cows, ‘P’ - wild boars. ‘PD’ - domesticated pigs. **b-d**. Quality metrics of Visium data, n = 7 subjects of 3 mammalian species examined in 3 independent experiments. Circles in c-d are medians, gray boxes span interquartile ranges. **e,f.** Boxplots showing the portal bias 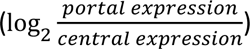 of secreted hepatocyte proteins (Methods) (e) and complement proteins (Methods) (f). P values are ranksum tests between humans and the other four species. Red lines are medians, boxes span interquartile ranges.

**Extended Data Figure 6.**
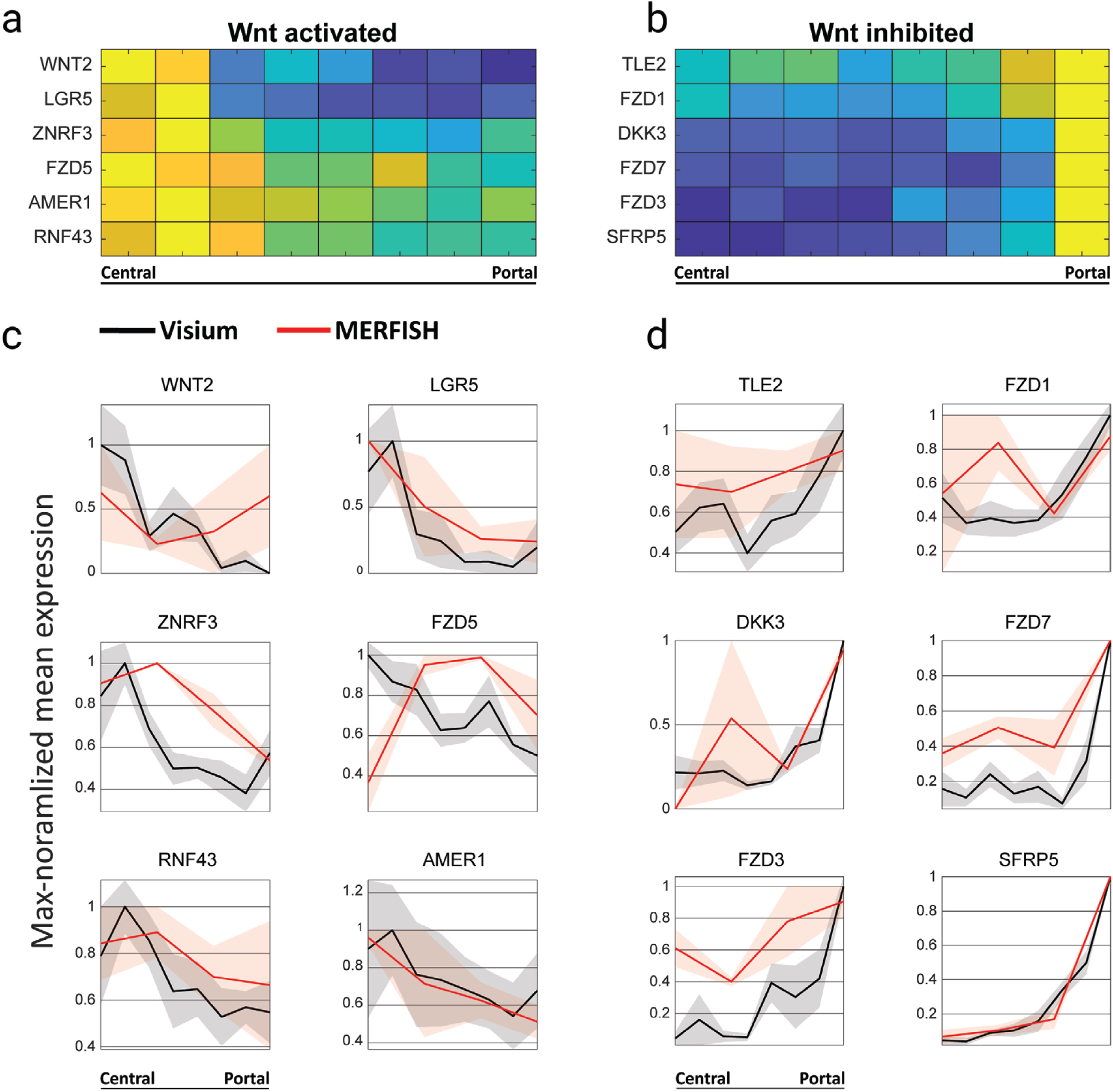
Zonation of Wnt pathway components. **a,b.** Heat maps for the Wnt-activated (a) and Wnt-inhibited highly (b) expressed genes (maximal expression above 1e-6). Shown are the six most pericentral and most periportal significantly zonated genes (qval<0.25) from the MERFISH Wnt gene sub-panel. Rows are the max-normalized mean expression of each gene based on the Visium data. (n = 5). **c.** Zonation profiles for the genes shown in (a). **d.** Zonation profiles for the genes shown in (b). Profiles in (c) and (d) are means over the patients normalized to their maximal expression level across zones; patches show the SEMs. MERFISH in red (n=2), Visium in black (n=5).

**Extended Data Figure 7.**
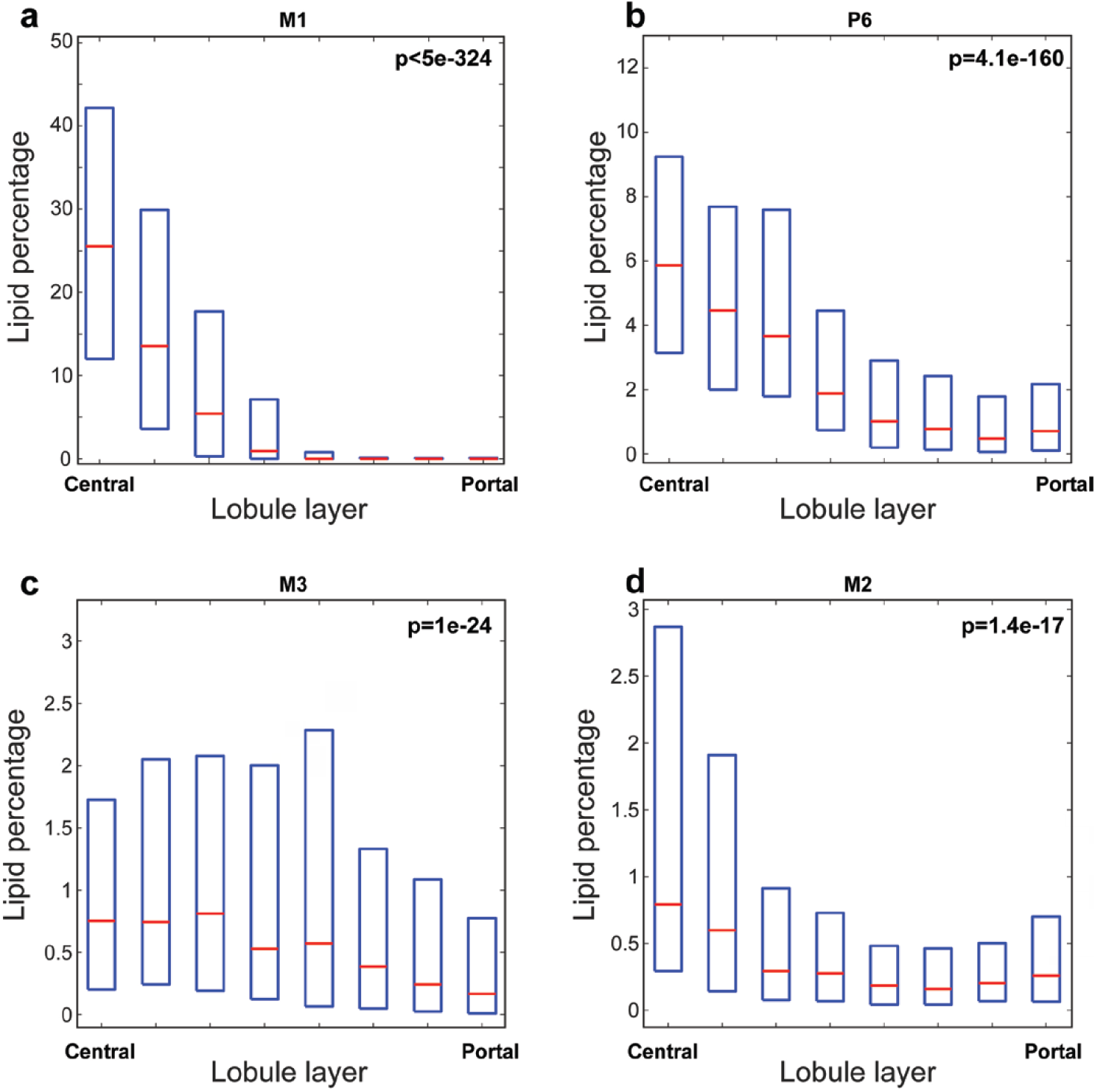
Lipid droplet enriched spots are pericentrally zonated. **a-d.** Boxplots showing the spots lipid content (in percentage from the spot’s area) along the porto-central axis. P values are Kruskal-Wallis tests between all zones. Red lines are medians, boxes span interquartile ranges.

**Supplementary Table 1** Pseudo-bulk gene expression levels from 16 Visium samples: Average gene expression levels calculated from all spots within the same patient (columns headings: PB (pseudo-bulk) and sample name).

**Supplementary table 2** Zonation table of healthy non-steatotic live liver donors’ samples (n=5, Fig. 2). The average zonal gene expression level along the porto-central axis. P value is calculated using Fisher’s combined probability test. Q value is calculated using the Benjamini-Hochberg procedure to control the false discovery rate. P values calculated for all genes that had maximal zonation level higher than 1e-6 in all the 5 patients.

**Supplementary Table 3** Gene expression table of the pericentral zone spots (layers 1 and 2) of liver samples of patients with mild steatosis along the gradient of lipid droplet percentile bins in the spot. (n=4, Fig. 4). Gradient of lipid droplet percentage was binned to 8 percentile bins between minimal (0%) and maximal spot lipid droplet content (89.6%). Average lipid-content bin gene expression levels were calculated. P-values were calculated using Kruskal Wallis tests, and q-values were determined using the Benjamini-Hochberg procedure to control the false discovery rate. K-means clustering (K=4) was performed on genes with a q-value threshold of 0.25, an expression level greater than 5e-6, and a dynamic range exceeding a 2-fold change in gene expression.

**Supplementary Table 4** Zonation tables all human liver samples (n=16, Fig. 1). The average zonal gene expression level along the porto-central axis. P values were calculated using Kruskal Wallis tests, and q-values were determined using the Benjamini-Hochberg procedure to control the false discovery rate.

**Supplementary Table 5** Zonation tables all non-human liver samples (n=7, Fig. 3). The average zonal gene expression level along the porto-central axis. SEM – Standard errors of the means.

**Supplementary Table 6** Spots lipid content percentage in the steatotic patients (n=4. M2, M3, M1, P6, Methods).

**Supplementary Table 7** Zonation tables of a healthy live human liver donor based on Visium HD (n=1, M6). The average zonal gene expression level along the porto-central axis. SEM – Standard errors of the means. P-values computed using Kruskal-Wallis method and Q-values computed using Benjamini and Hochberg correction for multiple hypotheses.

## Methods

### Experimental methods

#### Human sample preparation

Human samples and clinical data were obtained from patients undergoing either hepatectomy surgery due to a hepatic pathology, or a donor-hepatectomy for a living donor liver donation (Fig. 1b). In the hepatectomies that were performed due to liver pathologies, a sample was taken at least 5 cm from the closest margin of the pathology. In the hepatectomies that were performed as donor-hepatectomy for a living donor liver donation, a liver biopsy of 1 cm³ was obtained immediately upon abdominal opening, prior to mobilization of the liver and initiation of resection. All hepatectomies due to liver pathologies were performed at Sheba Medical Center, Israel with approval from the Sheba Medical Center Helsinki committee. All donor hepatectomies for a living donor liver donation were performed at Mayo-clinic, Rochester, Minnesota, with approval from the Mayo-clinic IRB committee. Informed consent was obtained from all patients, and all experiments followed all the Helsinki committee guidelines and regulations. Samples were taken only in cases where no pathology was observed in the sampling area during pre-operative and intraoperative evaluations. In all cases ∼1 cm of hepatic parenchyma distant from the liver capsule area were resected. Tissues were gently washed in PBS and were embedded in OCT for Visium and MERFISH or fresh-freezed for Visium HD.

#### Non-human sample preparation

Whole washed pig (*Sus scrofa domesticus*) livers were bought from a butcher at Kibbutz Lahav C.R.O., Israel, selling meat products from surplus animals. Specimens were adult males (5.5 months, ∼85kg) and an adult female (5.5 months, ∼75kg) raised under farming conditions. Washed cattle (*Bos taurus*) liver sections were donated by a butcher in Haifa, Israel. Specimens were adults males (17 months, ∼670kg) raised under farming conditions. Wild Boar (*Sus scrofa*) liver samples were taken from the cadavers of animals euthanized as part of population control regulated hunting done by the Israel Nature and Parks Authority (INPA). The sampling permit was acquired beforehand under INPA regulation. The researchers did not influence the time, location, or manner of death; they were solely permitted to take samples from the cadavers. In the methodological context of this study, the age and weight estimation of the specimens were conducted by trained INPA rangers and a veterinarian who participated in the hunting expedition. The boars’ ages were approximately 1-2 years, weighing approximately 60kg. Liver tissues were gently washed in PBS and were embedded in OCT for spatial transcriptomics (Fig. 3b-d, Extended Data Fig. 4).

#### 10x Visium spatial transcriptomics

Fresh frozen OCT embedded blocks were cryo-sectioned into 10µm thick slices and carefully placed on a Visium Spatial Gene Expression slide. Slides were fixed with pre-chilled methanol at-20 °C, stained for H&E as described in the Visium user guide and were captured using a Leica DMi8 widefield inverted microscope equipped with a Leica DFC7000T color camera and an HC PL APO 20x/0.80 DRY objective (506530, Leica Microsystems CMS GmbH). Permeabilization time was set according to the 10X Tissue optimization protocol, resulting in 24 min for all 16 human and 7 non-human mammalian samples. After tissue permeabilization and transcript capture, libraries were prepared according to the Visium Spatial Gene Expression User guide with 14-17 cycles of PCR for cDNA amplification. Four Libraries for each slide were pooled and loaded at a concentration of 400pM to a SP100 flow cell and sequenced on Illumina NovaSeq 6000. The Fastq reads generated from the sequencer were preprocessed by 10X Genomics space-ranger software (version 2.0.1) which included spatial de-barcoding, read-alignment to hg38 and UMI-generation. Further preprocessing was done in MATLAB R2022a.

#### 10x Visium HD Spatial Transcriptomics

Fresh-frozen tissue samples were processed into formalin-fixed, paraffin-embedded (FFPE) blocks. Sections of 5 μm thickness were mounted onto Visium HD slides and deparaffinized according to the Visium HD Spatial Gene Expression User Guide. Hematoxylin and eosin (H&E) staining was performed to visualize tissue morphology, and high-resolution brightfield images were captured using a Leica DMi8 widefield inverted microscope equipped with a Leica DFC7000T color camera and an HC PL APO 20x/0.80 DRY objective (506530, Leica Microsystems CMS GmbH). Following imaging, sections were destained and underwent decrosslinking as described in the Visium HD FFPE Tissue Preparation Handbook. Three specific probes for each targeted gene were added to the deparaffinized, destained, and decrosslinked tissues. Probe pairs hybridized to their complementary target RNA. Probe release and capture were performed using the Visium CytAssist instrument, followed by cDNA amplification and library construction. Our slide included sections from two patients – M2 and M6. The RNA yield of M2 was significantly lower (median of 83 vs. 169 UMIs per 8um*8um spot). We therefore continued this specific analysis with patient M6.

#### MERFISH spatial transcriptomics

Liver OCT blocks were sectioned to 10um sections. Sections were mounted on MERSCOPE glass slides (Vizgen) and were fixed with 4% PFA pre-warmed to 47 °C for 30 minutes, rinsed, permeabilized, and preserved in sterile 70% ethanol. The procedures of probe hybridization, gel-embedding, and tissue clearing were performed according to the MERSCOPE FFPE sample preparation user guide, using the predesigned pan cancer pathways 500 gene panel. Briefly, tissues were dried for 1h in room temperature to improve slide adherence. Then, tissues were photobleached for 3 hours using the Vizgen photo-bleacher, followed by a de-crosslinking step, gel embedding and tissue clearing for 3 days. Next, sections were incubated with MERFISH probe mix for 48 hours at 37 °C.

After probe hybridization, the samples were labeled with DAPI and poly T and placed in the MERSCOPE flow chamber for imaging. The raw image files obtained were processed via the MERlin image analysis pipeline^56^. The Cellpose software was utilized to segment cells from the nuclear DAPI signal. The decoded RNA molecules were then partitioned into individual cells to generate single cell count matrices.

### Computational Methods

#### Genome alignment

Human samples were aligned to genome assembly hg38. Domesticated pig samples were aligned to genome assembly Sscrofa11.1. Cattle samples were aligned to genome assembly BTauARS-UCD1.2. Boar samples were aligned to genome assembly Sscrofa11.1.

#### Spatial transcriptomics preprocessing

To remove outlier spots, for each patient, spots with log10 of total UMI counts lower than 3 standard deviations from the mean log10 of total UMI counts across all spots, or mitochondrial fraction over 5 standard deviations from mitochondrial fraction mean across all spots, were filtered out. Portal spots with prominent fibrosis, identified histologically based on the H&E staining, were manually marked and excluded from the analyses (Fig. 2a, patients M8, P7, P17, P2, P21). In addition, spots located in areas where the tissue anatomy integrity was unclear were manually annotated and filtered out. The remaining spots were used for analysis. To correct for high abundant transcript diffusion into neighboring spots^53^, we calculated for each gene the mean UMI count in spots within the fiducial frame that were not under the tissue to compute a background vector of expression. Next, the background vector was subtracted from each spot under the tissue, and negative values were set to zero. Finally, For the remaining spots, the raw data was normalized to relative counts per spot by dividing by the sum of all genes.

#### Pseudo-bulk analyses of human samples

Mitochondrial (^MT-) and ribosomal (^RPL, ^RPS) genes were removed from the variable gene list to minimize batch effect contribution to the analysis. Background subtracted counts were normalized, and a mean expression level was computed for each gene in every patient, creating a pseudo-bulk value for each gene. A Z-score was calculated for each gene across patients, and hierarchical clustering (Fig. 1c) and principal component analysis (Fig. 1d) were performed on the resulting Z-scores for all genes with maximal normalized expression above 5e-4. Spearman correlation distance and average linkage were used for hierarchical clustering. Comparison with NDD liver (Extended Data Fig. 1e) was performed on Visium data from Andrews et al.^9^. Pseudobulk was computed as the mean of spots annotated as hepatocytes and that further showed ALB expression above 0.003 of summed UMIs.

#### Human portal-central axis spot annotation

Visium: First, we extracted for each gene the spearman correlation coefficient between the gene’s expression level and *CYP2E1* expression level over all spots. The 20 genes that were most correlated or anticorrelated with *CYP2E1* over all spots were defined as our pericentral and periportal landmark gene sets (Pericentral landmark gene set: CYP3A4, ADH1B, CYP1A2, CYP2E1, APOA2, APOC1, ADH4, ADH1A, APOH, AMBP, GSTA2, ADH1C, SLCO1B3, AOX1, APOA5, DCXR, RBP4, OAT, CYP2C19, GC. Periportal landmark gene set: SERPINA1, APOA1, ALB, C7, NNMT, HAMP, ALDOB, ASS1, CYP2A7, MGP, A2M, FXYD2, CCL21, HAL, IGFBP2, SDS, AQP1, CYP2A6, FBLN1, PTGDS). Those landmark gene sets’ normalized summed expression levels enabled assigning a percentile-categorized zonation score to each spot (computed as the sum of periportal expression divided by the sum of periportal expression plus the sum of pericentral expression 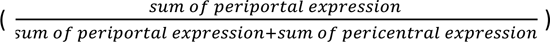. This zonation score was then binned into 8 zones with equal percentiles, and a zone index was assigned to each spot. To correct for spurious annotations, we applied a median filter by re-assigning the zone of each spot as the median zone of all spots surrounding it (relative distance of under 150µm, Fig. 2a). Subsequently, the zonation bias p-value of each gene was computed using the Kruskal Wallis test, followed by Benjamini and Hochberg correction for multiple hypotheses. Center of mass (COM) was calculated for every orthologous gene per sample as the weighted average of the zones^24,57^. Gene zonation scores were calculated as the linear combination of the zonal layer with the highest gene expression and the gene’s center of mass (Fig. 2, Extended Data Fig. 3). Gene zonation scores were non-defined for genes that exhibited low expression in our Visium datasets. For the MERFISH analysis, we used *NOTCH3* as a periportal landmark gene and reconstructed cell zonation using the distance from a *NOTCH3* positive cell. For the Visium HD analysis, we worked on the 8um spot size data. Non-parenchymal spots, identified histologically based on the H&E staining, were manually marked and excluded from the analyses, as were spots at the section periphery and spot with less than 50 UMIs (extended Data Fig. 3d). Zonation score was initially assigned to each spot using the same landmark gene method used for the Visium slides and binned into 8 zones based on equal percentiles over the analyzed spots. Subsequently, zone values were assigned the median zone of 100um*100um areas. Spots were binned and averaged, yielding the Visium HD-reconstructed zonation profiles.

#### Human hepatocyte marker gene Identification

To identify genes that serve as markers for hepatocytes, we utilized the single cell atlas generated by Massalha et al.^12^. We defined marker genes as those with a normalized expression level higher than 1e-5 and with a ratio of expression in hepatocytes to their maximum expression in all other cell types greater than 0.5. Since Hepatocytes contain more than 10-fold total mRNA than non-parenchymal liver cells, this value ensures that the majority of transcripts for such genes in each spot likely originate from hepatocytes^58^. We identified 1,724 hepatocyte marker genes (Fig. 2b,c, Fig. 3e).

#### Cross-species portal-central axis spot annotation

To remove outlier spots, for each non human sample spots with log10 of total UMI counts that are either higher or lower than 1.5 standard deviations from the mean log10 of total UMI counts across all spots were filtered out. For the non-human livers, each spot’s zone was determined based on landmark gene expression in cow and mouse^24^ and based on distances from the portal fibrotic septa in pigs and boars; Spots were then binned into 6 distinct zones, by equally proportioned percentiles and the assigned zone was median filtered using spots closer than 150µm. When comparing zonation trends of specific gene subsets (Fig. 3e, Extended Data Fig. 4e,f), the ratios between the mean expression in the two most periportal zones and the mean expression in the two most pericentral zones were computed. Only hepatocyte-specific genes that were highly expressed in at least 1 non-human species and in humans (max expression above 1e-5 in each) were analyzed.

#### Cross-species ortholog genes analysis

To allow for a common ground analysis, we sub-setted our data to genes that show 1:1 ortholog across all species (1 ortholog per gene per species). 11,345 Human, mouse, pig, boar, and cattle ortholog genes were found using the Ensembl biomart tool^55^. 1,344 hepatocyte-specific genes with orthologs across all species genes were found (Fig. 3e). Only human patients with steatosis level below 10% were used (n=7, M2, M3, M4, M5, M6, M7, M8). Secreted proteins were identified based on Zhang et al.^59^ as parsed in Halpern et al.^24^(Extended Data Fig. 4e), while complement proteins were derived from the KEGG^54^ Human Complement and Coagulation Cascade gene set (https://www.gsea-msigdb.org/gsea/msigdb/cards/KEGG_COMPLEMENT_AND_COAGULATION_CASCADES) (Extended Data Fig. 4f). For the analysis of Wnt-activated and Wnt-repressed genes we utilized the data from a published Wnt-hyperactivating liver-specific Apc knockout model mice^39^ and used the 20 genes that were most Wnt-up-regulated and the 20 genes that were most Wnt-down-regulated.

#### Lipid droplet quantification

Analysis of gene expression changes in early steatosis was performed on four out of the 7 patients that exhibited steatosis, and for which image quality was sufficient to reliably extract lipid content values (M1, M2, M3, P6). To extract the lipid percentage of each spot in the H&E attained Visium slide, an artificial neural network pixel classifier was trained on the H&E images using the QuPath software v.4.0^60^. The pixel classifier was trained to identify 3 histologic features: fat droplet, non-fatty liver tissue and the slide background. For each slide, the pixel classifier was trained on multiple fatty droplets and non-fatty tissue from the same H&E image (Fig. 4a), with the following parameters: Resolution: ‘very high’, channels: ‘Hematoxylin and Eosin’, Scales: ‘1,2,4’, Features: All 12 available features (the default is Gaussian). The spot locations were uploaded from the Visium output, and the QuPath measurement tools was used to quantify the number of fat droplet pixels and non-fatty liver tissue pixels in each spot (Fig. 4c).

## Data availability

Porto-central gene expression zonation profiles and gene expression changes along the steatosis gradient can be browsed using our web app (https://itzkovitzwebapps.weizmann.ac.il/webapps/home/session.html?app=HumanLiverZonation). Some browsers may need to refresh the page after loading the app. Spatial transcriptomics data of the human and non-human samples that were generated in this study can be downloaded from Zenodo, containing preprocessed 10X raw count tables, spot barcodes, tissue images and loupe browser files of all the human and non-human samples using the link https://zenodo.org/records/14795740?token=eyJhbGciOiJIUzUxMiJ9.eyJpZCI6Ijg5NzU1MzRjLTA2ZjYtNDU1ZS04MWZhLTRhNmIwODBkNDk4YSIsImRhdGEiOnt9LCJyYW5kb20iOiJhZDNmNDdjN2JlMjVmMTRhODZlMGU2NmI3NTE2NzA4NCJ9.5dbmnWYhg8UHjzXemALu5n94drsy3YBKgPALlqbwB1ZLrBEoIyw1vPJbU4cjV6IQS0Qd-QeXdaMfYtFdDXNCYg.

## Code availability

The code used to process the raw 10x Visium spatial transcriptomics data is available at https://github.com/OranYak/Human-liver. Any code for downstream analysis will be provided by the authors upon request.

## Author’s contributions

O.Y., A.A. and S.I. conceived the study. I.N., T.T., N.P. and R.P. operated on the patients and provided samples. O.Y., K.B.H., T.B. and Y.H. collected and processed human samples. A.A. and O.Y. collected and processed non-human samples. A.E. performed computational analysis. R.N. and Y.K.K. contributed to computational analysis. Visium experiments were performed under the guidance of M.K.,H.K., Y.H. and I,G., MERFISH experiments were performed under the guidance of L.F.A and D.H.. O.G. and Y.A. contributed to all microscopy imaging and pixel classification development. C.M. contributed to the histology slide evaluation. O.Y. and S.I performed computational analysis and wrote the manuscript. All the authors discussed the results and commented on the manuscript.

## Acknowledgements

The authors would like to thank R. Lapid, R. King, G. Kahila Bar-Gal, B. Rotblat, M. Zachut and J. Shpirer for their help in sourcing the non-human samples. S.I. is supported by the Helen and Martin Kimmel Award for Innovative Investigation, the Yad Abraham Research Center for Cancer Diagnostics and Therapy, the Moross Integrated Cancer Center, the Minerva Stiftung grant, a Weizmann-Sheba grant, the Israel Science Foundation grants no. 908/21 and 3663/21, the European Research Council (ERC) under the European Union’s Horizon 2020 research and innovation programme grant no. 768956 and a grant from the Ministry of Innovation, Science & Technology, Israel. Y.K.K. is supported by the JSMF Postdoctoral Fellowship in Understanding Dynamic and Multi-scale Systems. Illustration in Fig. 1a and mammals representative images in Fig.3 and Extended Data Fig. 5 were created using BioRender.

## References

1. Ben-Moshe, S. & Itzkovitz, S. Spatial heterogeneity in the mammalian liver. Nat Rev Gastroenterol Hepatol 16, 395– 410 (2019).

2. Gebhardt, R. Metabolic zonation of the liver: Regulation and implications for liver function. Pharmacol. Ther. 53, 275–354 (1992).

3. Cunningham, R. P. & Porat-Shliom, N. Liver Zonation - Revisiting Old Questions With New Technologies. Front Physiol 12, 732929 (2021).

4. Sahebjam, F. & Vierling, J. M. Autoimmune hepatitis. Front Med 9, 187–219 (2015).

5. Lindor, K. D. et al. Primary biliary cirrhosis. Hepatology 50, 291–308 (2009).

6. Afriat, A. et al. A spatiotemporally resolved single-cell atlas of the Plasmodium liver stage. Nature 611, 563–569 (2022).

7. Hildebrandt, F. et al. Host-pathogen interactions in the Plasmodium-infected mouse liver at spatial and single-cell resolution. Nat. Commun. 15, 7105 (2024).

8. Andrews, T. S. et al. Single-Cell, Single-Nucleus, and Spatial RNA Sequencing of the Human Liver Identifies Cholangiocyte and Mesenchymal Heterogeneity. Hepatol Commun 6, 821–840 (2022).

9. Andrews, T. S. et al. Single-cell, single-nucleus, and spatial transcriptomics characterization of the immunological landscape in the healthy and PSC human liver. J Hepatol 80, 730–743 (2024).

10. Aizarani, N. et al. A human liver cell atlas reveals heterogeneity and epithelial progenitors. Nature 572, 199–204 (2019).

11. Guilliams, M. et al. Spatial proteogenomics reveals distinct and evolutionarily conserved hepatic macrophage niches. Cell 185, 379–396.e38 (2022).

12. Massalha, H. et al. A single cell atlas of the human liver tumor microenvironment. Mol Syst Biol 16, e9682 (2020).

13. Brazovskaja, A. et al. Cell atlas of the regenerating human liver after portal vein embolization. Nat Commun 15, 5827 (2024).

14. Ramachandran, P. et al. Resolving the fibrotic niche of human liver cirrhosis at single-cell level. Nature 575, 512–518 (2019).

15. Matchett, K. P. et al. Multimodal decoding of human liver regeneration. Nature 630, 158–165 (2024).

16. Watson, B. et al. Spatial transcriptomics of healthy and fibrotic human liver at single-cell resolution. Preprint at 10.1101/2024.02.02.578633 (2024).

17. Prawira, A. et al. Single-cell and spatial atlas of steatotic liver disease-related hepatocellular carcinoma. Preprint at 10.1101/2024.05.09.593073 (2024).

18. Regev, A. et al. The Human Cell Atlas. Elife 6, (2017).

19. MacParland, S. A. et al. Single cell RNA sequencing of human liver reveals distinct intrahepatic macrophage populations. Nat Commun 9, 4383 (2018).

20. Hon, C. C., Shin, J. W., Carninci, P. & Stubbington, M. J. T. The Human Cell Atlas: Technical approaches and challenges. Brief Funct Genomics 17, 283–294 (2018).

21. Consortium, Gte. Human genomics. The Genotype-Tissue Expression (GTEx) pilot analysis: multitissue gene regulation in humans. Science 348, 648–60 (2015).

22. Hajaj, E., Pozzi, S. & Erez, A. From the Inside Out: Exposing the Roles of Urea Cycle Enzymes in Tumors and Their Micro and Macro Environments. Cold Spring Harb Perspect Med 14, (2024).

23. Chen, K. H., Boettiger, A. N., Moffitt, J. R., Wang, S. & Zhuang, X. Spatially resolved, highly multiplexed RNA profiling in single cells. Science 348, aaa6090 (2015).

24. Halpern, K. B. et al. Single-cell spatial reconstruction reveals global division of labour in the mammalian liver. Nature 542, 352–356 (2017).

25. Droin, C. et al. Space-time logic of liver gene expression at sub-lobular scale. Nat Metab 3, 43–58 (2021).

26. Ben-Moshe, S. et al. Spatial sorting enables comprehensive characterization of liver zonation. Nat Metab 1, 899–911 (2019).

27. Subramanian, A. et al. Gene set enrichment analysis: a knowledge-based approach for interpreting genome-wide expression profiles. Proc Natl Acad Sci U A 102, 15545–50 (2005).

28. Ramilowski, J. A. et al. A draft network of ligand-receptor-mediated multicellular signalling in human. Nat Commun 6, 7866 (2015).

29. Hu, S. & Monga, S. P. Wnt/-Catenin Signaling and Liver Regeneration: Circuit, Biology, and Opportunities. Gene Expr 20, 189–199 (2021).

30. Planas-Paz, L. et al. The RSPO-LGR4/5-ZNRF3/RNF43 module controls liver zonation and size. Nat Cell Biol 18, 467– 79 (2016).

31. Paris, J. & Henderson, N. C. Liver zonation, revisited. Hepatology 76, 1219–1230 (2022).

32. Forrest, A. R. et al. A promoter-level mammalian expression atlas. Nature 507, 462–70 (2014).

33. Chong, Z. X., Yong, C. Y., Ong, A. H. K., Yeap, S. K. & Ho, W. Y. Deciphering the roles of aryl hydrocarbon receptor (AHR) in regulating carcinogenesis. Toxicology 495, 153596 (2023).

34. Hildebrandt, F. et al. Spatial Transcriptomics to define transcriptional patterns of zonation and structural components in the mouse liver. Nat Commun 12, 7046 (2021).

35. Inoue, Y., Hayhurst, G. P., Inoue, J., Mori, M. & Gonzalez, F. J. Defective ureagenesis in mice carrying a liver-specific disruption of hepatocyte nuclear factor 4alpha (HNF4alpha). HNF4alpha regulates ornithine transcarbamylase in vivo. J Biol Chem 277, 25257–65 (2002).

36. Torre, C., Perret, C. & Colnot, S. Molecular determinants of liver zonation. Prog Mol Biol Transl Sci 97, 127–50 (2010).

37. Gougelet, A. & Colnot, S. A Complex Interplay between Wnt/β-Catenin Signalling and the Cell Cycle in the Adult Liver. Int J Hepatol 2012, 816125 (2012).

38. Burke, Z. D. & Tosh, D. The Wnt/beta-catenin pathway: master regulator of liver zonation? Bioessays 28, 1072–7 (2006).

39. Gougelet, A. et al. T-cell factor 4 and β-catenin chromatin occupancies pattern zonal liver metabolism in mice. Hepatology 59, 2344–57 (2014).

40. Thompson, M. D. & Monga, S. P. WNT/beta-catenin signaling in liver health and disease. Hepatology 45, 1298–305 (2007).

41. Matsumoto, S. & Kikuchi, A. Wnt/β-catenin signaling pathway in liver biology and tumorigenesis. Vitro Cell Dev Biol Anim 60, 466–481 (2024).

42. Wong, R. J. & Cheung, R. Trends in the Prevalence of Metabolic Dysfunction-Associated Fatty Liver Disease in the United States, 2011-2018. Clin Gastroenterol Hepatol 20, e610–e613 (2022).

43. Wong, V. W. et al. Impact of the New Definition of Metabolic Associated Fatty Liver Disease on the Epidemiology of the Disease. Clin Gastroenterol Hepatol 19, 2161–2171.e5 (2021).

44. Marra, F. & Svegliati-Baroni, G. Lipotoxicity and the gut-liver axis in NASH pathogenesis. J Hepatol 68, 280–295 (2018).

45. Matchett, K. P., Paris, J., Teichmann, S. A. & Henderson, N. C. Spatial genomics: mapping human steatotic liver disease. Nat. Rev. Gastroenterol. Hepatol. 21, 646–660 (2024).

46. Chalasani, N. et al. Relationship of steatosis grade and zonal location to histological features of steatohepatitis in adult patients with non-alcoholic fatty liver disease. J Hepatol 48, 829–34 (2008).

47. Brunt, E. M. Pathology of fatty liver disease. Mod. Pathol. 20, S40–S48 (2007).

48. Ishii, S., Iizuka, K., Miller, B. C. & Uyeda, K. Carbohydrate response element binding protein directly promotes lipogenic enzyme gene transcription. Proc Natl Acad Sci U A 101, 15597–602 (2004).

49. Roder, K., Zhang, L. & Schweizer, M. SREBP-1c mediates the retinoid-dependent increase in fatty acid synthase promoter activity in HepG2. FEBS Lett 581, 2715–20 (2007).

50. Altun, Ö. et al. Serum Angiopoietin-like peptide 4 levels in patients with hepatic steatosis. Cytokine 111, 496–499 (2018).

51. Utzschneider, K. M. & Kahn, S. E. The Role of Insulin Resistance in Nonalcoholic Fatty Liver Disease. J. Clin. Endocrinol. Metab. 91, 4753–4761 (2006).

52. Katz, J. & McGarry, J. D. The glucose paradox. Is glucose a substrate for liver metabolism? J. Clin. Invest. 74, 1901– 1909 (1984).

53. Guillot, A. et al. Mapping the hepatic immune landscape identifies monocytic macrophages as key drivers of steatohepatitis and cholangiopathy progression. Hepatology 78, 150–166 (2023).

54. Kanehisa, M. & Goto, S. KEGG: kyoto encyclopedia of genes and genomes. Nucleic Acids Res 28, 27–30 (2000).

55. Liberzon, A. et al. The Molecular Signatures Database Hallmark Gene Set Collection. Cell Syst. 1, 417–425 (2015).

56. Moffitt, J. R. et al. Molecular, spatial, and functional single-cell profiling of the hypothalamic preoptic region. Science 362, (2018).

57. Harnik, Y. et al. A spatial expression atlas of the adult human proximal small intestine. Nature 632, 1101–1109 (2024).

58. Halpern, K. B. et al. Paired-cell sequencing enables spatial gene expression mapping of liver endothelial cells. Nat Biotechnol 36, 962–970 (2018).

59. Zhang, Y. et al. Strategy for studying the liver secretome on the organ level. J Proteome Res 9, 1894–901 (2010).

60. Bankhead, P. et al. QuPath: Open source software for digital pathology image analysis. Sci Rep 7, 16878 (2017).

